# Using population-specific add-on polymorphisms to improve genotype imputation in underrepresented populations

**DOI:** 10.1101/2021.02.03.429542

**Authors:** Zhi Ming Xu, Sina Rüeger, Michaela Zwyer, Daniela Brites, Hellen Hiza, Miriam Reinhard, Sonia Borrell, Faima Isihaka, Hosiana Temba, Thomas Maroa, Rastard Naftari, Jerry Hella, Mohamed Sasamalo, Klaus Reither, Damien Portevin, Sebastien Gagneux, Jacques Fellay

## Abstract

Genome-wide association studies rely on the statistical inference of untyped variants, called imputation, to increase the coverage of genotyping arrays. However, the results are often suboptimal in populations underrepresented in existing reference panels and array designs, since the selected single nucleotide polymorphisms (SNPs) may fail to capture population-specific haplotype structures, hence the full extent of common genetic variation. Here, we propose to sequence the full genome of a small subset of an underrepresented study cohort to inform the selection of population-specific add-on SNPs, such that the remaining array-genotyped cohort could be more accurately imputed. Using a Tanzania-based cohort as a proof-of-concept, we demonstrate the validity of our approach by showing improvements in imputation accuracy after the addition of our designed addon SNPs to the base H3Africa array.

## 1. Introduction

By mapping the associations between single-nucleotide polymorphisms (SNPs) and various phenotypes, genome-wide association studies (GWAS) have allowed us to gain unprecedented knowledge on the genetic basis of various human diseases and traits. An important prerequisite to conducting GWAS is the availability of a cost-effective yet accurate high-throughput genotyping method. Genotyping arrays have been used widely over the past 15 years, including in many studies facilitated by biobank resources such as the UK Biobank[1]. However, genotyping arrays rely on the imputation of a sparse set of tag SNPs (e.g. millions of SNPs) to achieve acceptable density genome-wide (e.g. tens of millions of SNPs). The quality of imputation is dependent on the suitability of the tag SNPs and the similarity of haplotype structure between the reference panel and the study population[2, 3, 4, 5].

For study populations where a genetically similar reference panel or population-specific array content may not be available, whole-genome sequencing (WGS) offers an alternative to genotyping arrays. Previous studies have suggested that WGS may offer substantial gains in such a scenario, potentially pinpointing loci absent in GWAS conducted using genotyping arrays [6, 7]. However, due to the large sample sizes often required to gain sufficient statistical power in GWAS, the cost of WGS can still be prohibitive despite its recent decrease [8].

An alternative to WGS is the development of population-specific reference panels and genotyping arrays. For example, African-specific reference panels and genotyping arrays have been developed in recent years in an attempt to rectify the underrepresentation of African populations in genetic studies[9, 10, 11]. Notably, the Human Heredity and Health in Africa (H3Africa) consortium has developed the H3Africa genotyping array, which contains approximately 2.2 million tags, to capture genetic variability observed in various African populations [12]. Furthermore, the African Genome Resource (AFGR) reference panel has been designed to capture the haplotype structure of various African populations to improve imputation accuracy. However, driven by the long evolutionary history and lack of bottlenecks, the level of genetic diversity is much higher among African populations compared to non-African populations [13, 14]. Therefore, these resources have not yet been able to provide complete coverage of genetic variation across all African populations. For the remaining underrepresented populations, we propose the use of add-on SNPs as a cost-effective approach to improve genotype imputation.

In this paper, we present an approach to select population-specific add-on SNPs that supplement commercially available genotyping arrays. For a GWAS cohort, we propose to perform WGS in a small subset (e.g. 10% of the entire cohort), in order to supplement existing reference panels but also to inform the selection of the add-on content, such that the rest of the array-genotyped cohort could be more accurately imputed. Specifically, the WGS data could reveal population-specific allele frequency differences (Figure 1A and Figure 1B) and haplotype structure differences (Figure 1C). Such information enables the selection of add-on tag SNPs designed for the study population, such that the imputation of target SNPs that are poorly tagged by existing tag SNPs could be improved.

**Figure 1:**
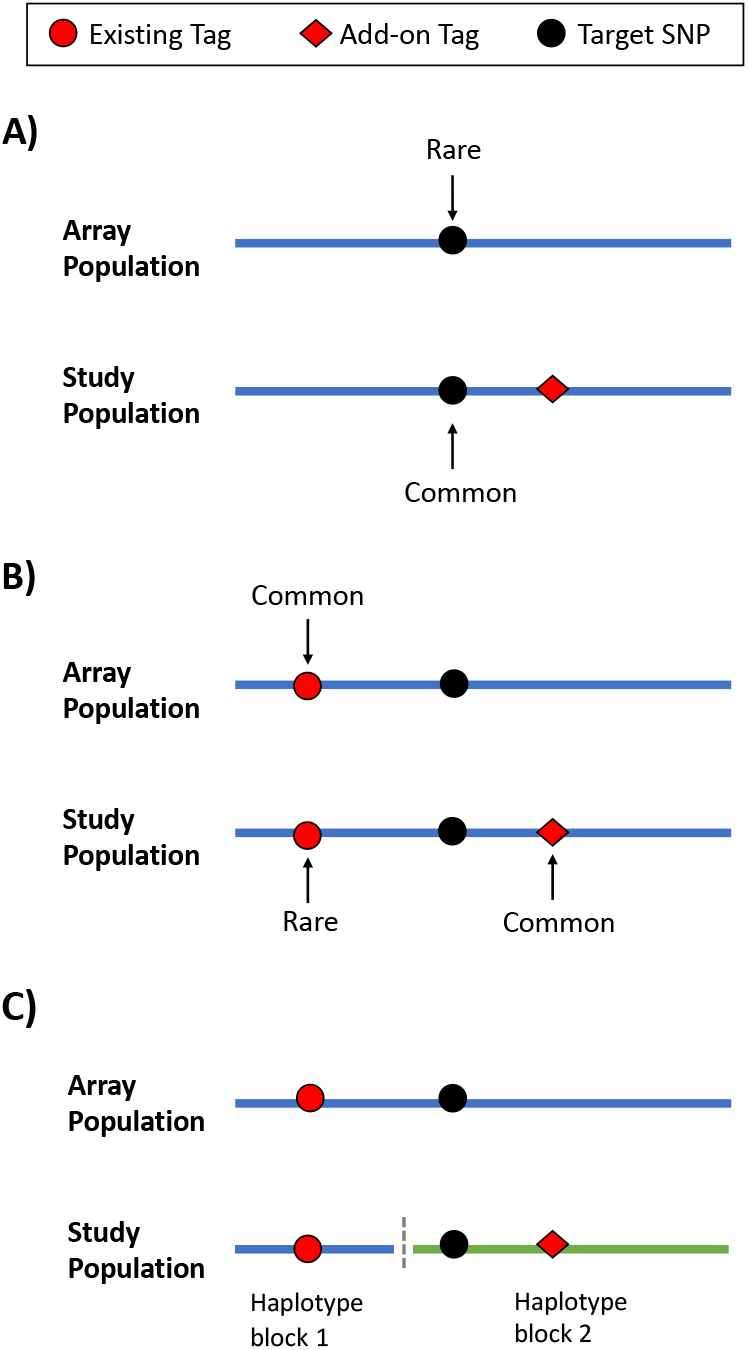
Scenarios under which add-on tags could improve genotype imputation. Array population represents the population that the existing genotyping array is designed for. Study population represents the population that the add-on tags are designed for. **A)** A target SNP that is rare in the array population, and was thus not designed to be tagged by any existing tag SNPs. However, it is common the study population, which justifies the use of an add-on tag. **B)** An existing tag SNP that is common in the design population but rare in the study population, thus reducing its tagging efficiency in the study population. **C)** The presence of population-specific haplotype structures in the study population, where the target SNP is no longer on the same haplotype block and no longer in strong LD with the existing tag SNP.

As a proof-of-concept example, we utilize 116 high coverage WGS samples from participants of the TB-DAR cohort (Tuberculosis patients recruited in a hospital in Dar es Salaam, Tanzania). Since the Tanzanian population is not incorporated in existing reference panels and array designs, including the AFGR reference panel and the H3Africa genotyping array, this cohort provides an ideal basis to evaluate our approach. We first illustrate the necessity for add-on SNPs by calculating the genetic differentiation between our Tanzanian cohort and other African populations. We proceed to select add-on SNPs that target common variants that are poorly imputed under the base H3Africa array. We then confirm the validity of our approach by evaluating the improvement in imputation accuracy enabled by the addition of add-on SNPs. Finally, we present an alternative selection scheme for mitochondrial and Y chromosome variants to improve haplogroup calling.

## 2. Material and Methods

### 2.1. Study description

This study was conducted based on a cohort of adult pulmonary tuberculosis (TB) patients from Dar es Salaam, Tanzania (TB-DAR). Participants were recruited at the Temeke Regional Hospital in Dar es Salaam. 128 patients were randomly selected from the cohort for WGS, and 116 samples which passed sequencing quality control were retained. Ethnic information of patients are based on self-reported information.

### 2.2. Whole genome sequencing and quality control

WGS was performed at the Health2030 Genome Center in Geneva on the Illumina NovaSeq 6000 instrument (Illumina Inc, San Diego CA, USA), starting from 1 μg of whole blood genomic DNA and using Illumina TruSeq DNA PCR-Free reagents for library preparation and the 150nt paired-end sequencing configuration. Average coverage was above 30 × for 75 samples, between 10× and 30× for 40 samples, and approximately 8× for a single sample.

Sequencing reads were aligned to the GRCh38 (GCA_000001405.15) reference genome using bwa[15] (Version 0.7.17), and duplicates marked using Picard (Version 2.8.14, http://broadinstitute.github.io/picard/). Following the GATK best practices (Germline short variant discovery)[16], Base Quality Score Re-calibration (BQSR) was applied using the GATK package[17] (Version 4.0.9.0). Variants were called individually per sample and then jointly. A Variant Quality Score Re-calibration (VQSR) based filter was then applied, with a truth sensitivity threshold of 99.7 and an excess heterozygosity threshold of 54.69. Samples with a high genotype missingness rate (> 0.5) were excluded.

To ensure that coordinates of the TB-DAR WGS data matched the GRCh37 based AFGR reference panel, a liftover was applied using Picard LiftoverVcf with the UCSC chain file (hg38ToHg19). Only SNPs that were successfully lifted over to the same chromo-some were retained. Within the X and Y chromosomes, SNPs within the pseudoautosomal regions[18, 19] were excluded.

### 2.3. Fixation index and genetic principal components

Relatedness between individuals within the TB-DAR WGS cohort and each African population of the 1000 Genomes project was calculated using KING[20]. Pairs up to first degree relatives were excluded.

To conduct principal component analysis (PCA), only autosomal SNPs that were genotyped in both 1000 Genomes and TB-DAR WGS cohorts were included. SNPs within long-range LD regions[21] were excluded. Using PLINK (Version 1.9)[22], LD pruning [23] (`plink --indep-pairwise 1000 50 0.05`) was applied and principal components were derived based on the merged cohorts (TB-DAR and all 1000 Genomes super-populations or TB-DAR and all 1000 Genomes African populations). To measure differentiation between the TB-DAR WGS cohort and various 1000 Genomes African populations, the fixation index (*F_ST_*) for each SNP was calculated using vcftools (v0.1.13)[24] according to Weir and Cockerham’s formulation [25]. Only autosomal SNPs that were genotyped and common (MAF> 0.05) in the merged cohort (TB-DAR and all 1000 Genomes African populations) were included. The reported genome-wide *F_ST_* measures were defined as the mean across the SNP-based *F_ST_* for all considered SNPs.

To estimate differentiation within a population, each population was divided into halves based on the median of the top genetic principal components. *F_ST_* was calculated between the two halves. Since the top genetic principal component explains the most proportion of genetic variability, this approach is expected to yield the two equally sized sub-populations that are the most differentiated within a population.

### 2.4. Selection of add-on SNPs

Our approach to select add-on SNPs can be divided into three main steps. In step 1, genotype imputation was performed. Poorly imputed SNPs were identified, and act as candidate target SNPs which our add-on tags would be designed to tag. In step 2, the optimal add-on tag SNPs were selected based on the population-specific LD structure and allele frequencies of the study cohort. In step 3, we evaluated the improvement in imputation performance when the selected add-on SNPs were incorporated onto the base H3Africa array. A summary of the approach can be found in Figure 2.

**Figure 2:**
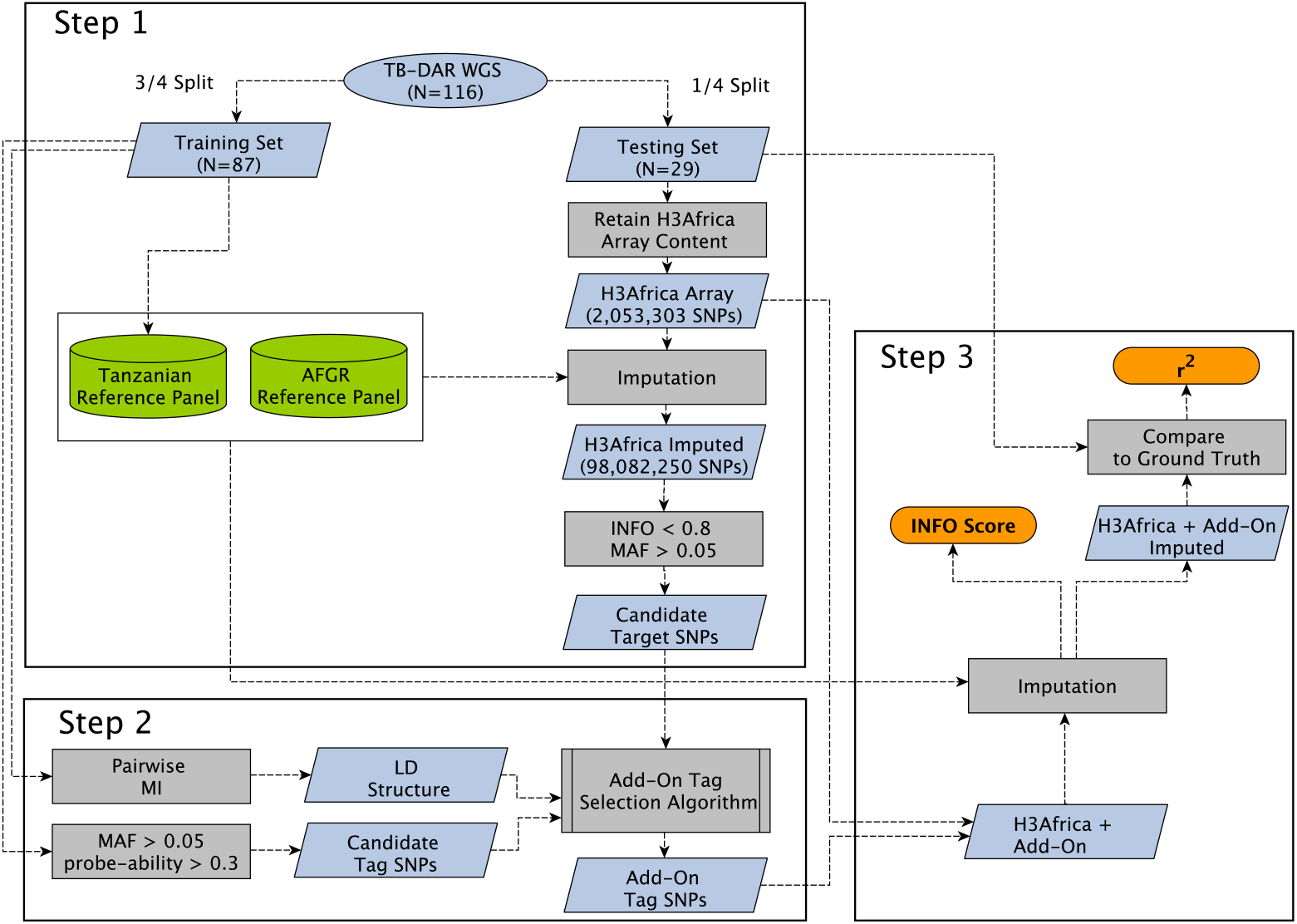
Schematic of our add-on tag SNP selection procedures, with steps illustrating: **Step 1)** Constructing a Tanzanian reference panel. Identifying candidate target SNPs, which are derived from poorly imputed SNPs when the H3Africa array is imputed based on the Tanzanian and AFGR reference panel. **Step 2)** Selecting add-on tag SNPs that optimally tag candidate target SNPs based on population-specific LD structures, allele frequencies, and probe qualities. **Step 3)** Evaluating improvements in imputation performance after adding add-on tag SNPs to the base H3Africa array. Calculating imputation quality metrics, including INFO score and r^2^ (correlation between imputed and sequencing-based genotypes). WGS, Whole-Genome Sequencing; AFGR, African Genome Resource; MAF,Minor Allele Frequency; MI, Mutual Information; LD, Linkage Disequilibrium.

#### 2.4.1. Step 1: Genotype imputation and identification of candidate target SNPs

The TB-DAR WGS cohort was divided into a training set (3/4 of the data) and a testing set (1/4 of the data).

To achieve optimal imputation accuracy, two reference panels were used to capture haplotype structures present in both the Tanzanian population and in other African populations. A custom Tanzanian reference panel based on the TB-DAR WGS training set samples was constructed using Minimac3[26]. The African Genome Resources (AFGR) reference panel (Web Resources) hosted on the Sanger imputation service (Web Resources)[27] was also utilized, where EAGLE2[28] was used for phasing and the positional Burrows-Wheeler transform (PBWT)[29] was used for imputation.

To identify poorly imputed SNPs expected under the H3Africa array content (Version 2, Web Resources), the TB-DAR WGS testing set was masked such that only SNPs present on the H3Africa array were retained. The masked data was imputed using both reference panels, and for each SNP the imputation was based on the reference panel that yielded a better imputation score. Candidate target SNPs were designated as SNPs that are poorly imputed (INFO < 0.8) but common in the TB-DAR WGS cohort (MAF > 0.05).

#### 2.4.2. Step 2: Add-on tag SNP selection

For each region, the set of candidate target SNPs (*S*_1_) was defined as SNPs that are poorly imputed but common (See Section 2.4.1). The set of candidate add-on tag SNPs (*S*_2_) was defined as sequenced SNPs that are common (MAF > 0.05), part of the AFGR Reference Panel or the TB-DAR reference panel, and available as Illumina Infinium probes (probe-ability score > 0.3). The set of existing tags (S3) was initialized as SNPs that are part of the H3Africa array.

LD information between SNPs were calculated based on TB-DAR WGS training set. We utilized mutual information (MI) as a LD metric (See Supplemental Methods), consistent with the choice of a previous array design study for the Japanese population [30].

To select the optimal set of add-on SNPs, we followed the framework of a forward-selection based algorithm [30]. In summary, the algorithm select tags that are in the strongest LD with the highest number of candidate target SNPs not captured by existing tags.

For a single iteration of the add-on tag SNP selection algorithm:

1. For a candidate target SNP (*j*), the existing tag SNP that is in strongest LD with it was identified. The MI score of the target SNP (*s_j_*) was defined as:

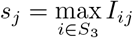

where *I_ij_* denotes the MI between SNP *i* and SNP *j.*
2. For each pair of candidate add-on tag SNP (*k*) and candidate target SNP (*j*), the add-on tag’s efficiency was defined as the expected change in MI (*δ_jk_*) resulting from the incorporation of the add-on tag:

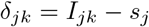
3. The efficiency of a candidate add-on tag SNP (*e_k_*) against all candidate target SNPs was defined based on the sum of the changes in MI:

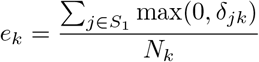

where *N_k_* denotes the number of probes required for the *k^th^* candidate add-on tag (2 for A/T or C/G SNPs, and 1 for all others).
4. The optimal add-on tag SNP (*k*\*) was identified based on the overall rank of its efficiency and probe-ability scores:

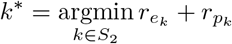

where *r_ek_* and *r_pk_* denotes the ranking of the efficiency score and probe-ability score respectively for the candidate add-on tag *k*.
5. *k*\* was added to the set of existing tags (*S*_3_), and the above steps were repeated. The selection procedure was stopped when there are no candidate add-on tags remaining (*S*_2_ becomes empty), or when the stopping criteria were met.

Figure S1 illustrates an example of a single iteration of the add-on tag SNP selection algorithm.

#### 2.4.3. Step 2: Region definitions and stopping criteria

To ensure the efficiency of add-on tag SNP selection but simultaneously guarantee sufficient coverage in prioritized regions, a two-step procedure for tag SNP selection with unique region definitions and stopping criteria was established.

Under Setting 1, regions spanning 5000 base pairs upstream and downstream of genes or SNPs associated with TB outcomes (reported by GWAS catalog [31], Open Targets[32], and other GWAS studies[33, 34, 35]) were considered. The killer cell immunoglobulin-like receptor (KIR) and human leukocyte antigen (HLA) gene regions were also considered. A region was subject to add-on tag SNP selection if it contained a substantial number of poorly imputed common SNPs, defined as more than 20% of SNPs with INFO < 0.8. Regions were also subjected to add-on tag SNP selection if it contained an uneven spatial distribution of well imputed common SNPs, defined as the spread of poorly imputed SNPs (INFO < 0.8) being more than 1.25 times the spread of well-imputed SNPs (INFO ? 0.8). To guarantee sufficient coverage, iterations of the forward-selection algorithm was run for each region independently until less than 0.5% of candidate target SNPs within the region showed *δ_k_* improvements. The process was then repeated for each of the prioritized regions.

Under Setting 2, the selection of add-on tag SNPs was expanded to any region across the genome that contained poorly imputed common SNPs. The regions were defined as either a haplotype block (`plink --blocks`)[36, 22] or a region spanning 5000 base pairs upstream and downstream a candidate target SNP, whichever larger. To maximize the selected add-on tag SNPs’ tagging efficiencies, a single iteration of the algorithm was run concurrently across all regions. The tag SNP that scored the best across all regions was incorporated. The process was then repeated until the total number of budgeted add-on probes (N=5000) has been exhausted.

#### 2.4.4. Step 3: Evaluation of imputation accuracy

The TB-DAR WGS testing set was utilized to measure improvements in imputation performance enabled by the add-on tag SNPs. For all target SNPs tagged by at least one add-on SNP, imputation quality (INFO score) derived from the base H3Africa array was compared against imputation quality derived from the H3Africa array with the addition of add-on tags. In addition, to measure the accuracy of the imputed genotypes, squared Pearson correlation coefficients (r^2^) was calculated between the imputed genotype dosages (0,1 or 2) and the ground truth dosages based on the WGS data.

### 2.5. Y Chromosome and Mitochondrial Haplogroups

The haplogroups of TB-DAR participants were called using HaploGrep2[37] and yhaplo[38] for the mitochondria and the Y chromosome respectively. The Phylotree mitochondrial[39] and Y chromosome[40] phylogeny databases were used to identify marker SNPs. Marker SNPs for each main haplogroup that any TB-DAR participant was part of were included as add-on SNPs, if not already existing on the H3Africa array. In addition, we added maker SNPs 2 branch points below the main haplogroup that any TB-DAR participant was part of.

## 3. Results

### 3.1. Differentiation between the Tanzanian population and other African populations

Study participants of the TB-DAR WGS cohort originated from various ethnic groups within Tanzania (Table S1). A majority of participants belonged to the Bantu-speaking ethnic groups (*N* = 108, 93.1%), with a small minority that belonged to the Nilotic (*N* = 1, 0.8%) and Cushitic (N = 3, 2.6%) speaking ethnic groups. Self-reported ethnic information was not available for four participants.

To quantify the population differentiation between the TB-DAR WGS cohort and the 1000 Genomes African populations, for each pair of populations we calculated the genome-wide fixation index (*F_ST_*). Figure 3A illustrate the pairwise *F_ST_* measures between the TB-DAR WGS cohort and 1000 Genomes African population, along with their respective sampling locations. In general, genetic differentiation was greater between populations that are further away geographically. For example, TB-DAR displayed the least differentiation with the Bantu-speaking Luhya population (LWK) in neighbouring Kenya, but the most differentiation with West African populations such as the Gambian in the Western Division of Gambia (GWD) and the Mende in Sierra Leone (MSL). A similar pattern was observed among 1000 Genomes African populations (Figure 3B), where population pairs in the same geographic region (e.g., YRI and ESN) were among the least differentiated population pairs. In addition, the genetic principal components (PCs) shown in Figure S2 also illustrate a similar pattern, where distances in PC space approximately scaled with geographic distances between the sampling locations of populations.

**Figure 3:**
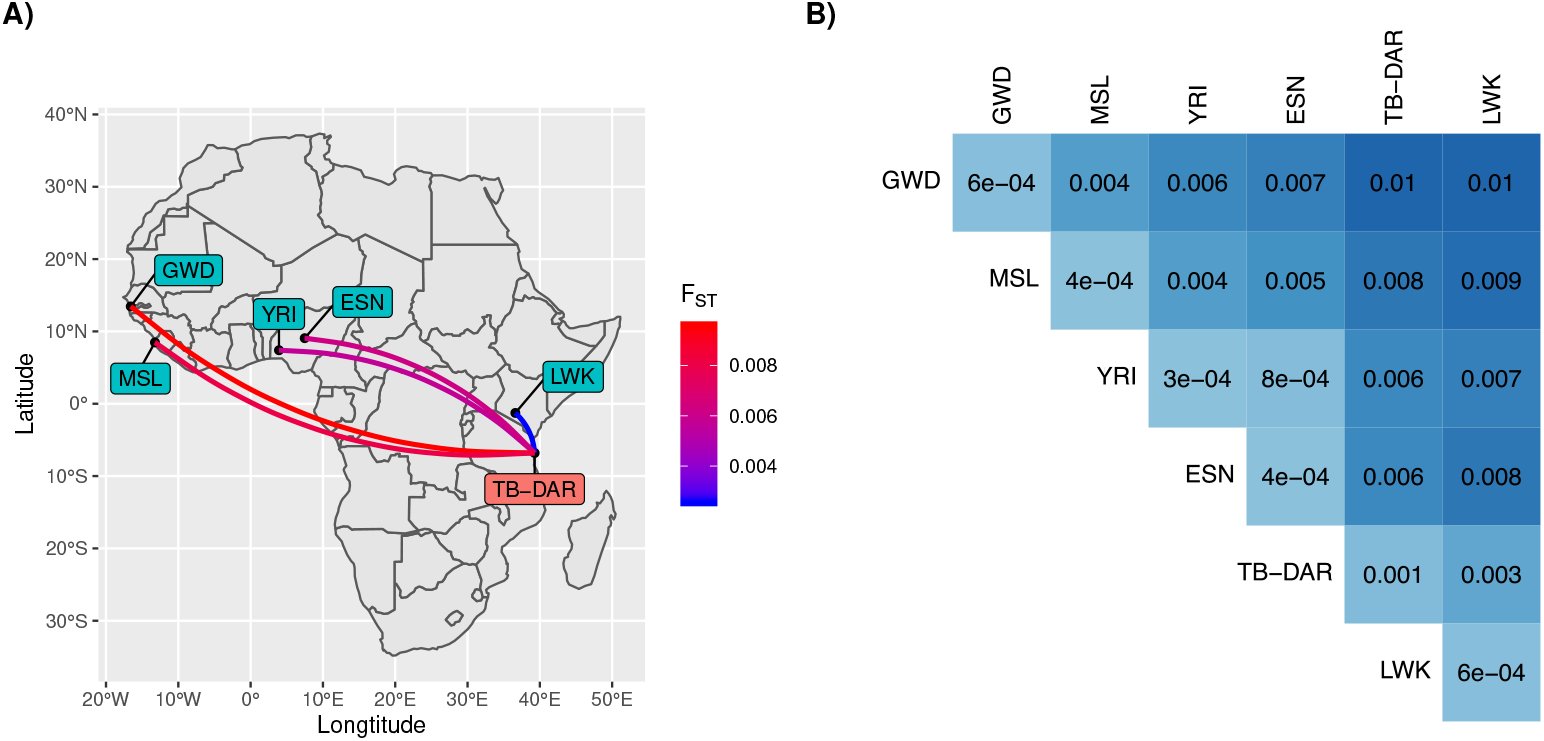
Genetic differentiation of African populations **A)** Sampling locations of 1000 Genomes African populations and the TB-DAR WGS cohort. Line colors illustrate the degree of differentiation (*F_ST_*) between TB-DAR and 1000 Genomes populations. **B)** Pairwise *F_ST_* measures between 1000 Genomes African population and TB-DAR. Diagonals of the matrix represent differentiation within a population, calculated between two halves of the population defined based on the median of the top genetic principal component. 1000 Genome Populations: GWD - Gambian in Western Divisions in the Gambia; MSL - Mende in Sierra Leone; YRI - Yoruba in Ibadan, Nigeria; ESN - Esan in Nigeria; LWK - Luhya in Webuye, Kenya.

To further evaluate the significance of differentiation between populations, we compared the inter-population *F_ST_* against the within-population *F_ST_*. The within-population *F_ST_* was calculated between two halves of each population that are expected to be the most differentiated, defined based on the median of the top genetic principal component. The diagonal of Figure 3B represent within-population *F_ST_* measures. For every population, the within-population *F_ST_* was lower than the inter-population *F_ST_* against the population which it is the least differentiated from. For example, the within-population *F_ST_* of the TB-DAR WGS cohort (0.001) is lower than the inter-population *F_ST_* against the LWK population (0.003).

These results quantify the genetic diversity of populations within Africa, and illustrate the differentiation between the TB-DAR cohort and African populations of the 1000 Genomes project. Thus, the need to supplement external reference panels with Tanzanian specific haplotypes and to design population-specific add-ons for the TB-DAR cohort is warranted.

### 3.2. Selection of add-on SNPs and improvements in imputation accuracy

The selection of add-on SNPs was conducted under two different settings (Section 2.4.3). Under a coverage-guaranteeing setting (Setting 1), we selected 1669 add-on SNPs within 337 prioritized TB-associated regions. In addition, under an efficiency-driven setting (Setting 2), we selected 2734 further add-on SNPs across the rest of the genome. Figure S3 shows the distribution of all selected SNPs across chromosomes.

To confirm the validity of our approach, we used the TB-DAR WGS testing set to compare the imputation accuracy based on the base H3Africa array against the improved H3Africa array with our add-on content. Figure 4A shows the mean imputation quality of target SNPs that our add-on SNPs were designed to tag across different minor allele frequency (MAF) percentile bins. Under both settings, we observed strong overall improvement across MAF bins in imputation accuracy with the incorporation of add-on tag SNPs, reflected by the increase in mean INFO score and *r*^2^ (correlation with WGS ground truth). While the magnitude of increase in mean imputation accuracy was similar for both settings, in general, target SNPs in prioritized regions were better imputed. This was as intended since, under Setting 1, even relatively well-imputed SNPs within each region would be tagged by add-on SNPs in order to guarantee coverage.

**Figure 4:**
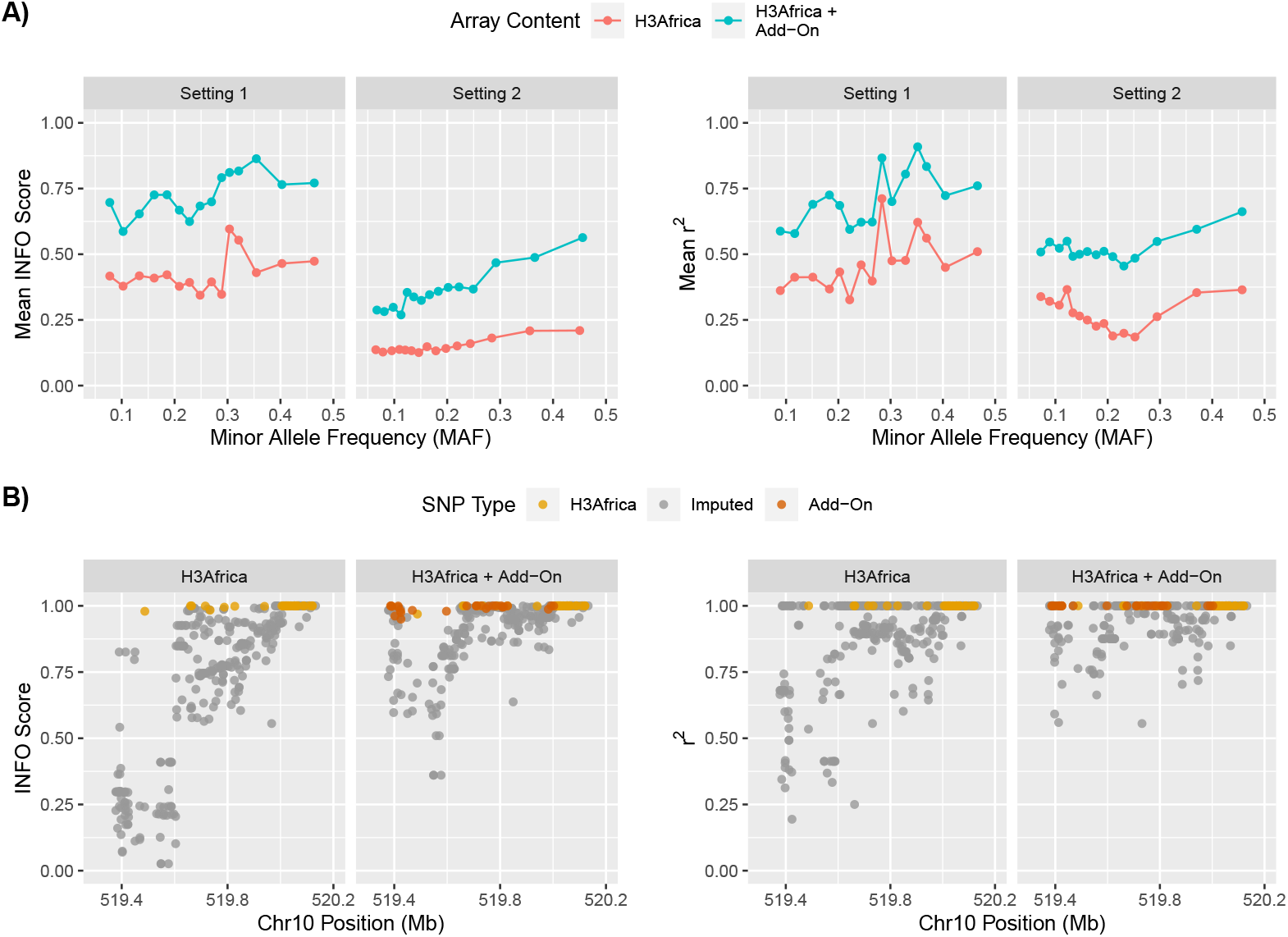
Improvement in imputation performance subsequent to the addition of add-on tag SNPs. **A)** Mean INFO score and *r*^2^ (between imputed and sequenced ground truth) of target SNPs designed to be tagged by add-on SNPs, prior and subsequent to the incorporation of add-on SNPs. Facet grids illustrate results based on two tag SNP selection settings: coverage guaranteeing within prioritized regions (Setting 1) and efficiency driven in all other regions (Setting 2). **B)** Example region on chromosome 10 where the incorporation of add-on tag SNPs lead to the increase in imputation performance. Facet grids illustrate imputation performance prior and subsequent to the incorporation of add-on tags. Color of dots represent type of SNP (existing H3Africa tags, add-on tags, or any other imputed SNPs).

An example region where our approach functioned as expected is shown in Figure 4B. Our designed add-on SNPs lead to improved imputation of target SNPs, reflected by increases in both INFO score and *r*^2^. Noticeably, add-on SNPs were mainly located in proximity to the previously poorly imputed target SNPs (left side of the region). This indicates, as designed, that only add-on SNPs that are in relatively strong LD with target SNPs were selected, as LD generally scales inversely with distance.

To quantify the efficiency of the selected add-on SNPs, Table 1 shows the number of targeted SNPs with INFO score improvements. Under an INFO score threshold of 0.8 (commonly used in GWAS), our 4403 add-on SNPs would allow the incorporation of an additional 10,349 and 38,336 target SNPs in GWAS, in TB associated regions (Setting 1) and all other regions (Setting 2) respectively. This translates to the addition of approximately 6 and 14 target SNPs per add-on SNP, under Setting 1 and Setting 2 respectively. As expected, the number of successfully tagged target SNPs per add-on SNP is lower under Setting 1. This is because to guarantee coverage, relatively short haplotypes are tagged, resulting in the reduced efficiency of each add-on tag SNP.

**Table 1:** Efficiencies of add-on tag SNPs, categorized based on source reference panel and selection settings. Imputation improvements categorized as any increase in INFO score, or any increase that resulted in INFO score exceeding 0.8 when previously under 0.8.

| Reference Panel | Tags Added | INFO Score Improvement | INFO Score > 0.8 |
| --- | --- | --- | --- |
| **Prioritized Regions (Setting 1) - Coverage Guaranteeing** |  |  |  |
| All | 1669 | 52,798 | 10,349 |
| Tanzanian | 666 | 33,516 | 2881 |
| African Genome Resource (AFGR) | 1003 | 20,010 | 6753 |
| **Other Regions (Setting 2) - Efficiency Driven** |  |  |  |
| All | 2734 | 26,3192 | 38,336 |
| Tanzanian | 1417 | 16,7040 | 8749 |
| African Genome Resource (AFGR) | 1317 | 95,892 | 26,564 |

### 3.3. Mitochondria and Y chromosome haplogroups

Since mitochondrial and Y chromosome haplogroups provide an efficient manner to track human evolutionary history, we targeted haplogroup markers to improve the accuracy of haplogroup calling. The distribution of mitochondrial and Y chromosome haplogroups within the TB-DAR WGS cohort are shown in Figure S4A and Figure S4B respectively. With regards to the mitochondrial DNA, most individuals belonged to the L haplogroup. This was consistent with findings based on the 1000 Genomes project[41], where the L haplogroups were found to be the dominant haplogroups in African pop-ulations. For the Y chromosome, a majority of male individuals belonged to the E haplogroups, with a small minority belonging to the B, R, and others. This was also consistent with the 1000 Genomes project[42], where the E haplogroups were found to be dominant in African populations. Also in the Luhya population in neighbouring Kenya a small minority belonged to the B haplogroup[42].

To ensure that our add-on content includes haplogroup markers that complement the existing content on the H3Africa array, we selected 103 and 31 haplogroup marker SNPs as add-ons for the mitochondria and Y chromosome respectively. For the mitochondria, we saw an average improvement in haplogroup calling of 22% compared to the H3Africa array.

For the Y chromosome, due to the limited number of add-on SNPs and sufficient coverage by the H3Africa array, we did not observe any significant differences in haplogroup calling.

## 4. Discussion

The strategy to supplement external reference panels with WGS samples from an internal study cohort has been employed by previous studies[43, 44]. Specifically, it has been shown that the addition of even a relatively small number of samples from the internal cohort leads to improved imputation accuracy, especially if the study population is genetically dissimilar from the populations captured by existing reference panels[45, 6]. Our work confirms the utility of including population-specific haplotypes in the reference panel used for imputation, but it also shows that the use of add-on SNPs further improves imputation accuracy of common variants in the study population.

Our add-on tag SNP selection procedure did not explicitly target population-specific SNPs, such as ancestry informative markers[46, 47], but rather targeted any SNP observed in our study population that are expected to be poorly imputed under the existing base array content. Such a choice was driven by the aim of GWAS, which is to map any SNP associated with the trait of interest, which may not necessarily be population-specific. Nevertheless, we did apply an allele frequency based (MAF) cutoff to ensure that only SNPs polymorphic in the study population were targeted. As a result, a substantial fraction of the targeted SNPs were successfully imputed based on the TB-DAR reference panel (Table 1). This suggested that our add-on SNPs were able to tag population-specific haplotype structures, which contributed to improved imputation accuracy.

An add-on tag SNP that most efficiently tags a target SNP (in the strongest LD) may not necessarily be the optimal tag, as the genotyping error rate of the probe for the particular SNP may be high. To rectify such issue, we limited our selection to add-on tags SNPs with probes that have high success rates (Illumina probe-ability score > 0.3), and weighted the trade-off between LD strength and probe quality equally when selecting the optimal add-ons. Nevertheless, a more complex weighting scheme may result in even better performance.

We introduced two settings for the selection of add-on SNPs, namely either coverage-guaranteeing (Setting 1) or efficiency-driven (Setting 2). For users of our approach, the number of regions assigned to each setting could be adjusted depending on the study. For example, if there exists strong prior knowledge with regards to genes implicated in or loci associated with the trait of interest, these regions could be assigned to Setting 1. Conversely, for traits with a lack of prior knowledge, a greater proportion of regions could be assigned to Setting 2, such that tag selection would be conducted in a more hypothesis-free manner.

A limitation of our approach is that only common SNPs (MAF > 0.05) were targeted by the selected add-on SNPs. Such a choice was made due to the limited sample size of our WGS cohort, where for rarer target SNPs there would be insufficient observations to estimate LD. Nevertheless, the imputation accuracy of rarer SNPs (for example, 0.01 < MAF < 0.05) which are in strong LD with the targeted SNPs could still increase if tested in a larger testing set.

In conclusion, in order to improve imputation accuracy in populations underrepresented in existing reference panels and genotyping array designs, we propose a framework where a subset of a cohort is sequenced and the rest genotyped using an array supplemented with the selected add-on SNPs. Using a Tanzanian-based cohort as a proof-of-concept, we demonstrated that under our approach, the WGS data could be leveraged to supplement existing reference panels and to select add-on SNPs, such that imputation accuracy is improved. Our approach is generalizable to any other population to improve genotype imputation, and thus provides a cost-effective solution to increase the power of GWAS in a diverse range of underrepresented populations and to further our understanding of human genetic diversity.

## Supplemental Data

Supplemental Data include 5 figures and 1 table.

## Ethics Approval and Consent

Ethical approval for the TB-DAR cohort has been obtained from the Ethikkomission Nordwest-und Zentralschweiz, the Ifakara Health Institute and the National Institute for Medical Research in Tanzania. An informed consent has been obtained from every patient who has been recruited into the TB-DAR cohort. This consent includes the use of the patient’s blood for human genomic analyses.

## Acknowledgments

This work was supported by the Swiss National Science Foundation (Sinergia Grant: 177163). We thank the study participants of the TB-DAR study for their contribution. We thank K. Harshman, I. Bartha, C. Howald, and D. Lamparter (Health 2030 Genome Center, Geneva, Switzerland) for sequencing support. We thank C. Thorball, K. Popadin, D. Lawless, O. Naret, and F. Hodel for helpful discussion.

## Data and Code Availability

Software code and a list of add-on SNPs designed for the TB-DAR cohort is available at: https://github.com/zmx21/h3africa-addon

## Web Resources

Open Targets: https://www.targetvalidation.org/

GWAS Catalog: https://www.ebi.ac.uk/gwas/

Sanger Imputation Service: https://imputation.sanger.ac.uk/

AFGR Reference Panel: https://imputation.sanger.ac.uk/?about=1#referencepanels H3Africa Genotyping Array: https://chipinfo.h3abionet.org/help

