## Supplemental Tables and Figures for "Using population-specific add-on polymorphisms to improve genotype imputation in underrepresented populations"

### Supplemental Data

#### *A1. Supplemental Tables*

Table S1: Self-reported ethnic groups of study participants of the TB-DAR WGS cohort, categorized by major linguistic groups.

| Ethnic Group | N |
|---|---|
| **Bantu (N=108)** | |
| Chaga | 3 |
| Digo | 1 |
| Fipa | 1 |
| Gogo | 4 |
| Ha | 4 |
| Haya | 1 |
| Hehe | 2 |
| Jita | 2 |
| Kwaya | 1 |
| Kwere | 1 |
| Luguru | 2 |
| Makonde | 6 |
| Makua | 3 |
| Matumbi | 3 |
| Mwera | 5 |
| Ndamba | 1 |
| Ndengereko | 11 |
| Ngindo | 3 |
| Ngoni | 2 |
| Nyakyusa | 2 |
| Nyamwezi | 3 |
| Nyaturu | 1 |
| Nyiramba | 2 |
| Pare | 3 |
| Pogolo | 5 |
| Sambaa | 8 |
| Sukuma | 1 |
| Yao | 2 |
| Zaramo | 21 |
| Zigua | 4 |
| **Nilotic (N=1)** | |
| Assa | 1 |
| **Cushitic (N=3)** | |
| Iraqw | 1 |
| Rangi | 2 |
| **Not Reported (N=4)** | |

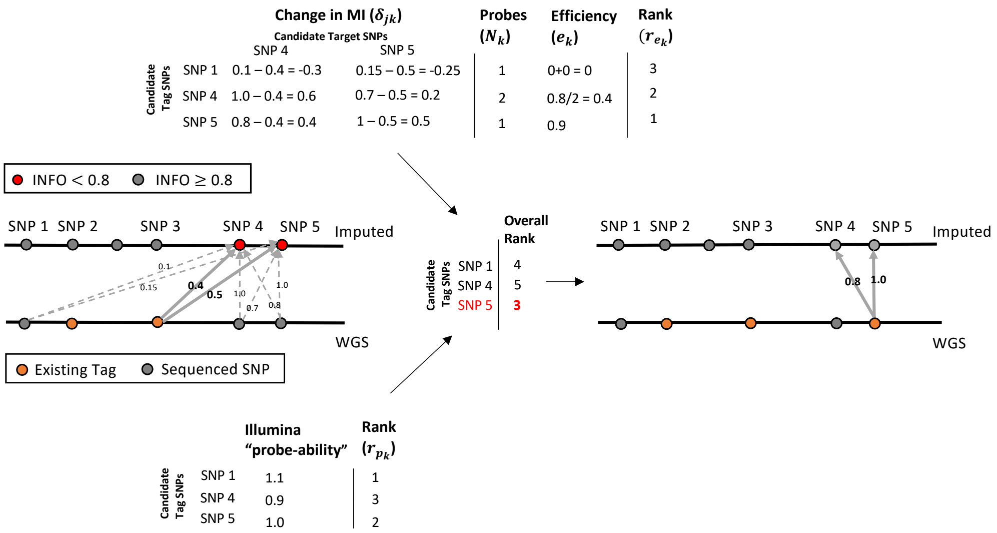


Figure S1: Schematic illustrating a single iteration of the add-on tag SNP selection algorithm. Edges between SNPs represent the strength of Linkage Disequilibrium (LD), measured by Mutual Information (MI). The optimal candidate tag SNP is selected based on the best overall rank, taking into account the efficiency ($e_k$), the number of probes required ($N_k$), and the quality of the probe (Illumina probe-ability). In subsequent iterations, the newly added tags are Incorporated as existing tags, such that the change in MI ($\delta_{jk}$) incorporate the contribution of add-on tags.

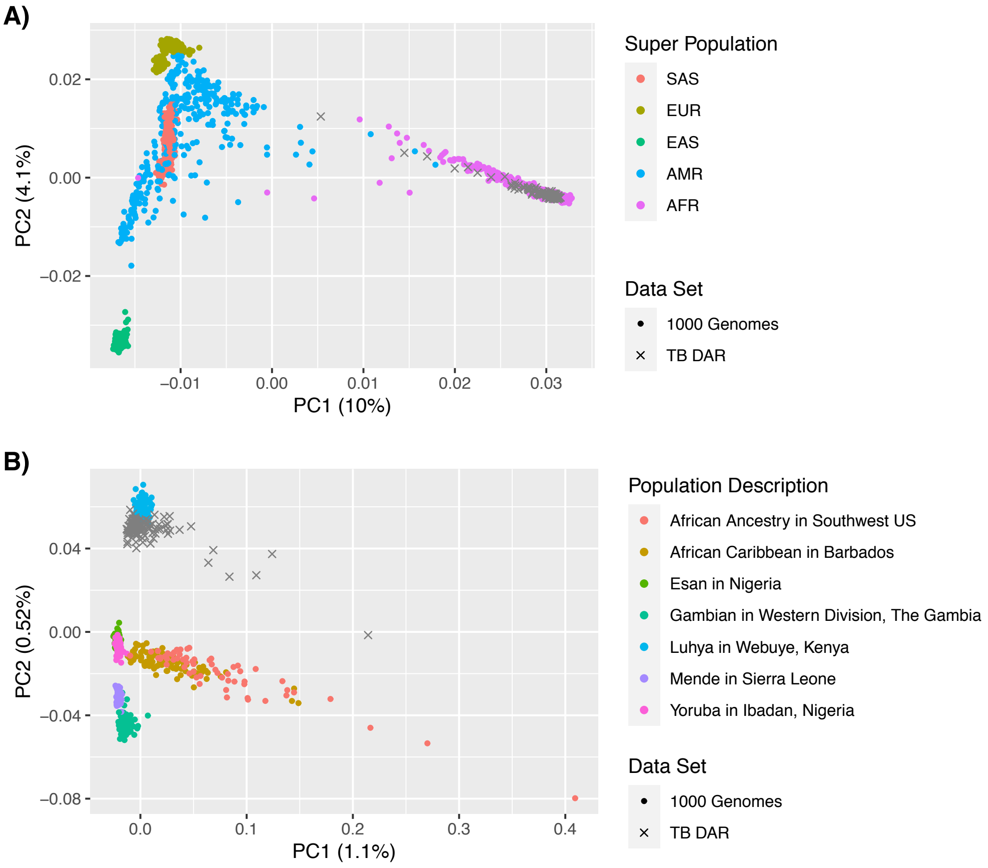


Figure S2: Genetic principal components (PCs) based on SNPs genotyped in both the TB-DAR WGS and 1000 Genomes cohort. Percent of variance explained by each PC are indicated in brackets. **A)** All 1000 Genomes populations, grouped according to super-populations. **B)** 1000 Genomes African populations.

1000 Genomes Superpopulations: SAS, South Asian; EUR, European; EAS, East Asian; AMR, Ad Mixed American; AFR, African.

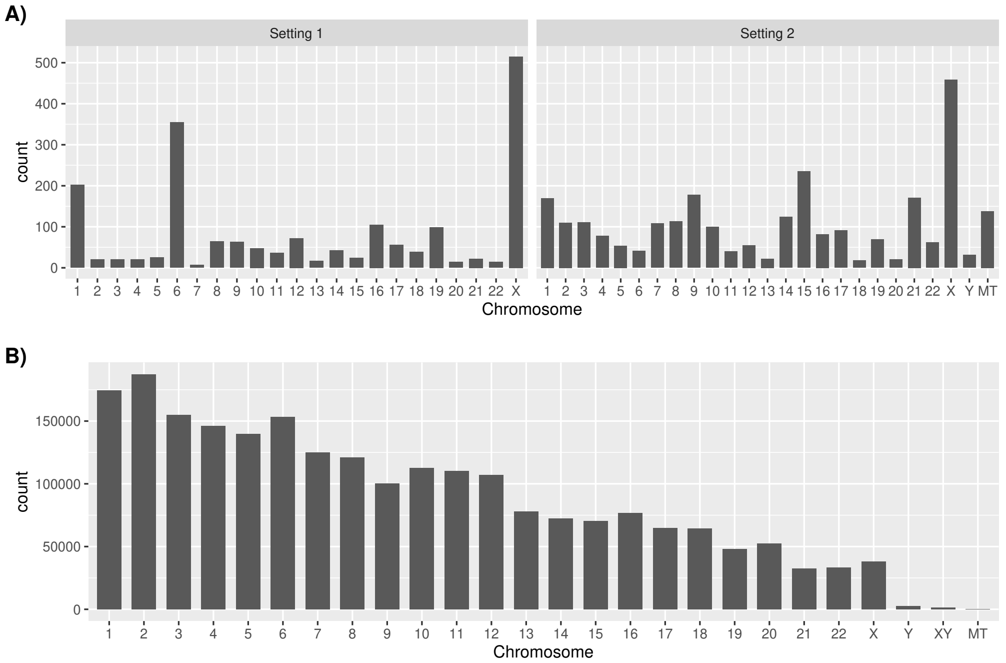


Figure S3: Number of tag SNPs on each chromosome and the mitochondria. **A)** Add-on tag SNPs selected based on Settting 1 and Setting 2. **B)** Existing tag SNPs on the H3Africa array.

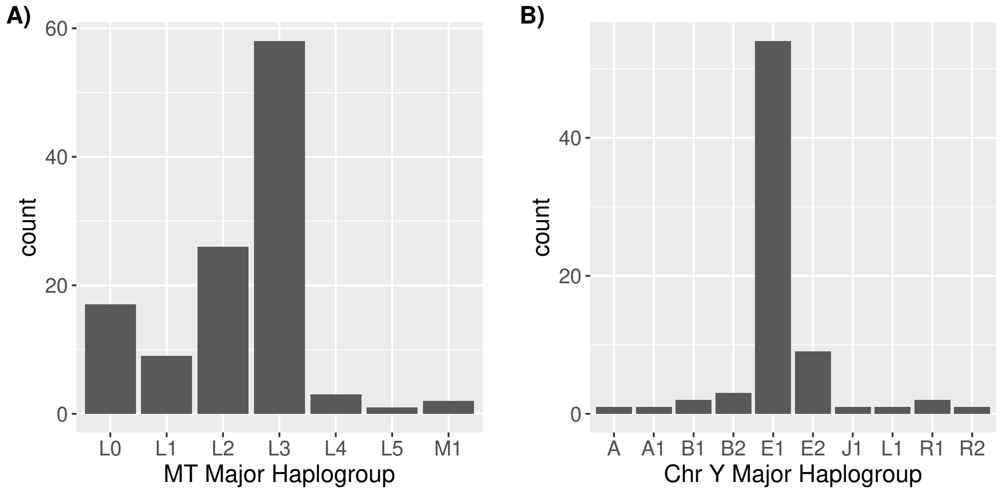


Figure S4: Haplogroups of participants within the TB-DAR WGS cohort for the **A)** Mitochondria **B)** Y chromosome (Males Only).

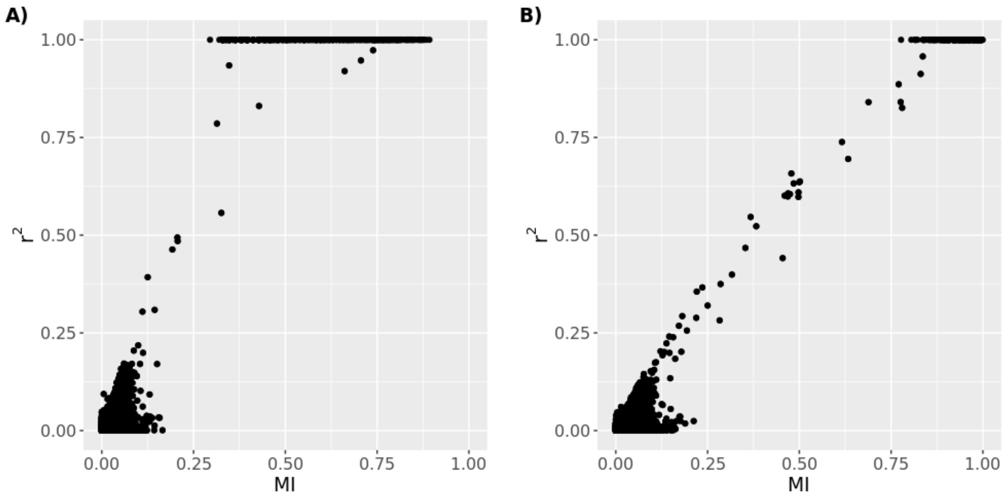


Figure S5: Comparison of two LD metrics: pairwise $r^2$ and Mutual Information (MI). LD was measured between 500 random pairs of SNPs. **A)** SNPs with low minor allele frequencies ($0.05 \leq$ MAF $\leq 0.25$) **B)** SNPs with high minor allele frequencies ($0.25 \leq$ MAF $\leq 0.5$).

*A3. Supplemental Methods*

To measure the pairwise LD between a candidate tag SNP and a candidate target SNP, we utilized pairwise mutual information (MI). The pairwise MI between SNP $i$ and SNP $j$ ($I_{ij}$) was defined as:

$$I_{ij} = H_i + H_j + H_{ij}$$

where $H_i$ denotes the entropy of a single SNP and $H_{ij}$ denotes the joint entropy defined as:

$$H_i = -\sum_{k=0}^{2} \frac{N(g_i = k)}{N} \log_2 \left( \frac{N(g_i = k)}{N} \right)$$

$$H_{ij} = -\sum_{k=0}^{2} \sum_{k'=0}^{2} \frac{N(g_i = k, g_j = k^{'})}{N} \log_2 \left( \frac{N(g_i = k, g_j = k^{'})}{N} \right)$$

where $N(g_i = k)$ denotes the number of individuals with genotype $k$ for SNP $i$, $N(g_i = k, g_j = k^{'})$ denotes number of individuals with genotype $k$ for the SNP $i$ and genotype $k^{'}$ for the SNP $j$, and $N$ denotes the total number of individuals.

Figure S5 illustrates that the differences between MI and $r^2$ (classical measure of LD). Pairwise MI was weaker than $r^2$ between random pairs of lower frequency SNPs ($0.05 \leq$MAF$\leq 0.25$), but similar to $r^2$ for higher frequency SNPs ($0.25 \leq$MAF$\leq 0.5$). This indicates that $r^2$ tend to overestimate the correlation between lower frequency SNPs. Thus, MI was chosen to indirectly favour the inclusion of higher frequency SNPs as add-on tag SNPs.
